# Association between movement patterns, microbiome diversity, and potential pathogen presence in free-ranging feral pigeons foraging in dairy farms

**DOI:** 10.1101/2023.10.11.561861

**Authors:** Miranda Crafton, Shai Cahani, Avishai Lublin, Luise Rauer, Orr Spiegel

## Abstract

The feedback between host behavior and disease transmission is well acknowledged, but empirical studies demonstrating associations between individual’s pathogens or microbiota composition and their movement are rare. We investigated these associations in feral pigeons (*Columba livia domestica*), a synanthrope species known to host a plethora of zoonotic pathogens. We captured pigeons in three dairy farms along an urbanization gradient in central Israel. We combined GPS-tracking with Next Generation Sequencing to characterize pigeons’ movement and microbiota, respectively. We found that pigeons roosted primarily in human settlements, with frequent visits to dairy farms and other agricultural sites. Microbiota diversity and composition varied between sites and the individuals within them, and several pathogens relevant to poultry, cattle, and human-health were frequently detected. Pigeons in the urban site covered shorter distances and carried a greater diversity of bacteria compared to those in rural sites. Intriguingly, beyond these among-site differences, exploratory individuals (measured by the number of unique locations they visited) had more diverse microbiota. We conclude that pigeons can potentially serve as transmission vectors among wildlife, livestock, and humans . Further, the associations between host behavior and microbiota diversity emphasize the relevance of wildlife movement analyses for disease ecology and One Health.

## Introduction

Pathogen transmission typically requires the movement of hosts or vectors between locations, highlighting the fundamental role of animal movement in understanding dynamics for the diversity of both pathogenic (and non-pathogenic) microbes across large and small scales (Dougherty et al. 2017). Behavioral feedback between host and parasites further stresses the inherent connection between movement and disease (Ezenwa et al. 2016). Host movement determines the exposure rate to different pathogens, as well as their potential to transmit them into new sites. Explorer individuals (that move more across the habitat), for instance, are likely to have more connections in their social network with consequences for transmission. The pathogens in turn are not only affected by the host movement but may affect it either indirectly (e.g. reducing movement due to reduced body conditions; (Fofana and Hurford 2019), or directly through deliberate behavioral manipulations (Poulin and Maure 2015). Accordingly, the fields of animal movement and disease ecology have drawn much attention from scientists looking to investigate the relationship between host movement patterns and their role in disease transmission.

Zoonotic diseases (pathogens that jump species barriers from animals to humans) are an emerging issue because the interactions among wildlife, livestock, and humans are intensifying due to the population growth of the latter two (Craft 2015). A recent review identified 175 animal-borne pathogens that are associated with emerging infectious diseases (Woolhouse et al. 2001), and Covid19 is a current and urgent reminder of the devastating and lasting effects of zoonotic outbreaks (Mackenzie and Smith 2020). Zoonotic diseases are estimated to account for 15.8% of all deaths around the world (and up to 43.7% in low-resource countries), with more than 2.7 million human deaths per year (Eubank et al. 2004; Salyer et al. 2017). To optimize global health, the One Health approach was created with the goal of using integrated knowledge across animals (both wildlife and livestock), humans, and their environment (Rabinowitz et al. 2013; Davis et al. 2017). Nevertheless, while One Health literature has often focused on broad ecological scales to study the origin, transfer, and potential vectors of zoonotic pathogens in an attempt to predict and prevent outbreaks, mechanistic examples focusing on transmission and feedbacks in specific systems are often left out.

Bio-logging techniques, such as GPS telemetry, may provide a detailed picture of host movement, bridging these knowledge gaps and identifying where, when, and how individuals move over time for predicting the pathways of potential diseases (Daversa et al. 2017). Accumulating knowledge in the field demonstrates how movement patterns emerge from complex interactions between individual needs (e.g. to find food or shelter), capacities (e.g. the flight capacity), and external conditions (e.g. resource distribution; (Nathan et al. 2008). Indeed, species show a remarkable variation across sites, and among individuals, directly affecting their transmission potential (Dougherty et al. 2017, Spiegel et al. 2017).

Similarly to the bio-logging improvements, ongoing technological improvement in high-throughput DNA sequencing (also known as Next Generation Sequencing) provides important insights into transmission dynamics of both pathogenic and non-pathogenic microbes between hosts, and the influence of host behavior on the microbiome. Gut microbiome is affected by host environment (Loo et al. 2019), lifestyle, social interactions and behavior (Sarkar et al. 2020). For example, natural bird movement has been linked to increased microbiome diversity (Corl et al. 2020), and food variation between different foraging sites has been shown to influence bird microbial composition (Gadau et al. 2019). The microbiome can serve as a pathway that connects an individual’s environment to health disparities, and differences in microbial composition across populations have important implications for the functions of the microbiome and eventually host health (Kuthyar and Reese 2021). For example, one study found significant correlation between microbiome composition and infection of Avian Influenza Virus, where infection status dramatically influenced microbial community variation in five different species of ducks (Hird et al. 2018). Understanding such interactions and relationships enables the use of innovative and holistic approaches to diagnosis, treatment, and intervention (Trinh et al. 2018).

Agricultural sites and practices are a common environmental factor in exposing humans and livestock to wildlife populations, and the diseases they may carry. Dairy and poultry farming are particularly known to attract a variety of wildlife from peripheral habitats, due to their resource availability including food, shelter, and possible security from natural predators who avoid human habitats. European starlings (*Sturnus vulgaris*) for example, often aggregate in high numbers from surrounding cities and towns to forage at farms, creating an intersection for otherwise distant subpopulations (Cabe 2021). Such use of shared space together with high densities and mobility may facilitate pathogen transmissions from birds to cattle or birds to other birds. Considering contacts with workers (at the farm) and members of the community (at urban roosts) who regularly interact with these wildlife and domestic species highlights the potential harmful epidemiological consequences of these encounters between humans, animals, and the environment.

Indeed, birds have received a lot of attention in the field of zoonotic diseases due to their mobility and potential to spread pathogens over large distances and ecological barriers (Nabi et al. 2021, Cabe 2021). While large-scale movements such as migration offer important insights into disease transmission, routine local movements within populations (e.g. foraging between resource patches) are also crucial for transmission dynamics. Perhaps the most suitable model system for investigating the association between movement, microbiome diversity and pathogen transmission is the ubiquitous feral pigeon (*Columba livia domestica;* hereafter simply pigeon). Pigeons are common synanthropes found in both urban and agricultural environments worldwide, often sharing their space with humans (Mia et al. 2022). They are social animals, aggregating into large flocks at resource-rich locations, or roosts at human buildings. Their home range sizes are determined by the availability and distribution of preferred resources, resulting in spatial overlap between various surrounding subpopulations (Haag-Wackernagel and Moch 2004, Nabi et al. 2021). They are known reservoirs for pathogens and carry a variety of agricultural and public health-relevant viral, bacterial, fungal, and parasitic pathogens; a comprehensive review by Haag Wackernagel and Moch in 2003 found that feral pigeons have been documented to carry at least 60 different human pathogenic organisms such as Salmonella, Chlamydia, Cryptococcus, and Escherichia as some examples. Among these 60 diseases, at least seven have been reportedly contracted by humans (Osman et al. 2013). Between 1941 and 2004 there have been at least 207 documented transmissions of disease-causing pathogens from pigeons (Haag-Wackernagel and Moch 2004). Indeed, environments shared by dairies, humans, and pigeons are known to increase the possibility of spillover among host species, demonstrating pigeons’ importance in the context of disease transmission (Elser et al. 2019).

Only a handful of studies investigated pigeon movement (Carlson et al. 2011, Sol and Senar 1995) and microbiome diversity (Grond et al. 2019) despite their relevance for host-parasite feedbacks. Here we address this knowledge gap and study these two components at three dairy farms in central Israel. First we ask: do pigeons at these farms differ in their movement, and do individuals within each site differ from each other? Previous studies have found that groups of pigeons who occasionally forage together may actually roost separately, and diverge in their movement (Rose et al. 2006), thus likely resulting in differential exposure levels and transmission potential. Each of our three study sites differ in level of urbanization, and we expect that individuals at the more urban site (Mikve Israel) will move less than the other two sites (Maale Hahamisha and Mevo Horon) due to proximity to surrounding food availability and roosting sites (Tucker et al. 2018). Second, we ask which pathogens do pigeons carry at our focal dairy farms? Here we expect a diverse list of bacteria (including cattle, and human-relevant ones). This expectation reflects the above-mentioned literature regarding pigeon’s diverse pathogenic loads, as well as a couple of local studies reporting *Salmonella* (Yeruham et al. 2006; Osman et al. 2013) and *E Coli* (Vasconcelos et al. 2018). Finally, we ask if individual movement is associated with microbial diversity? The microbiome is influenced by a variety of environmental factors such as habitat, diet, and more (Amato et al. 2015, Hyde Embriette R. et al. 2016). Yet, how movement correlates with microbiome is poorly studied. In a rare example, Corl et al (2020) reported that alpha diversity in barn owls’ microbiome was higher in individuals that moved greater distances away from the nest each day. Such a positive association can reflect, for instance, exposure to more diverse bacterial environments. Hence, we predict that exploratory individuals who visited more sites throughout their daily foraging activity would display more diverse microbiota. To answer these questions we combined pigeon GPS tracking data and untargeted next-generation RNA sequencing and taxonomic profiling of a subset of the individuals.

## Methods

### Study Sites

The fieldwork for this project took place at three different dairy farms in central Israel representing a gradient of increasing urbanization (**Figure 1A,B)**: Mevo Horon, Maale Hahamisha, and Mikve Israel. Mevo Horon (31.8496° N, 35.0350° E, +250 masl) is a settlement located 13 km northwest of Jerusalem. It is very isolated, with almost no other human settlements within a 5km vicinity, and no urbanized (built) areas within 2km area around this settlement (see Methods section in Supplementary Materials for quantification procedure from areal imaging). Kibbutz Maale Hahamisha is located on a high elevated ridge (31°49.239N, 35°6.667E, +800 masl) about 7km southeast of Mevo Horon. This site is surrounded by approximately 4 small human settlements within ∼2km distance (covering around 13.3% of the area) and several others within a 5 km range. Lastly, Mikve Israel (32.0312° N, 34.7833° E, +30 masl) is located approximately 30 km further northwest from Mevo Horon, within the heavily populated coastal plane. It is at the outskirts of the Tel Aviv metropolitan, surrounded by immediate urban environments and additionally home to poultry houses nearby the dairy farm. Accordingly, 42% of the land surrounding this site is urban, making it our most urban site.

**Figure 1.**
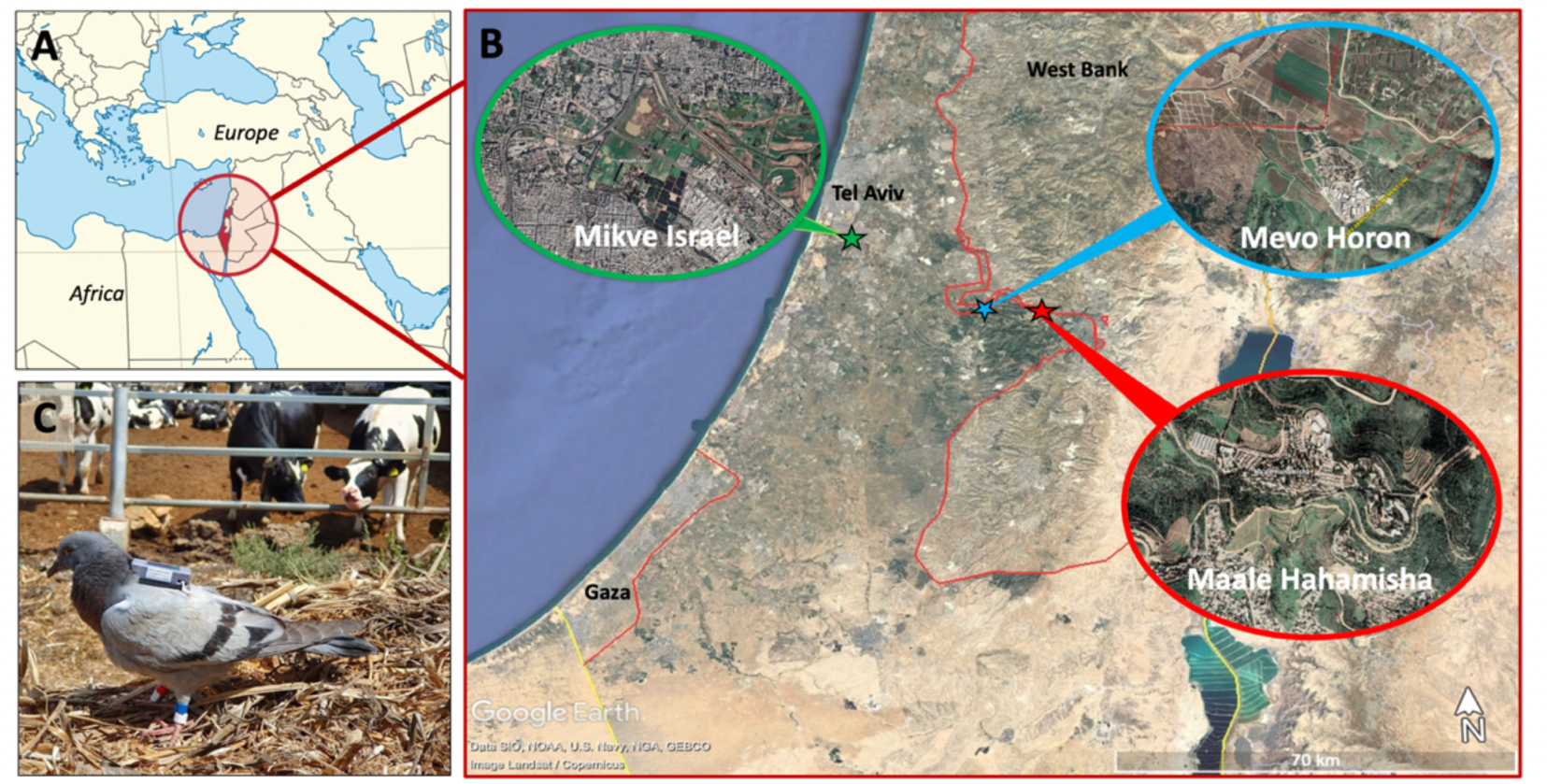
The dairy farm - feral pigeon (Columba livia domestica) study system. (**A**) A map of the Mediterranean and Israel’s position (**B**). A focal map of the three study sites within Israel; Mevo Horon being the most rural, Maale Hahamisha is intermediately rural, and Mikve Israel is urban. (**C**). Pigeon tagging and banding. Color-bands enable individual identification in the field. A subset of individuals are equipped with GPS-GSM transmitters in a backpack configuration. Photo credit: O. Spiegel.

### Pigeon Captures, GPS-Tagging, and Microbiome Sampling

We captured a total of 308 pigeons using a combination of mist nets, walk-in ground traps, and falling traps with grains spread nearby to lure individuals towards the traps. Captures took place between July 2019 to November 2021 throughout all seasons. After capture, morphological measurements were recorded from each pigeon following standard ornithology procedures for weight, age (juveniles/adults), wing and tail length, color, and chest feathers collection for sexing (by https://www.karnieli-vet.com/). All pigeons were fitted with metal rings containing a unique ID, in addition to an individual-specific combination of colored rings.

Captured individuals that were determined as adults in good body condition (and above the 250g minimal weight threshold) were fitted with an Ornitela OrniTrack 10g GPS transmitter attached in a backpack configuration (**Figure 1C**). These tags employ state-of-the-art technology, including solar power and data download capacity over GSM communication, allowing extended tracking duration. To save energy tags used a diurnal duty cycle (dawn to dusk) reflecting pigeons’ activity. During day time, pigeon locations were sampled at 10 minute interval when the battery was properly charged (reducing fix rate to once an hour or once a day if battery was below 50% or 25% power, respectively). Examples of a typical track after filtering can be seen in the appendix (**Figure S1**). In total, 44 individual pigeons were fitted with GPS tags (Maale Hahamisha: 24, Mevo Horon: 11, Mikve Israel: 9).

To determine microbiome composition, we swabbed all captured individuals for the collection of DNA samples. We first swabbed the choanal cleft, followed by the cloaca. We chose this combined swabbing method due to our interest primarily in the overall microbial composition of each individual, rather than place of occurrence within an individual. The swab was then placed into 2mL of Viral Transport Medium (BHI) containing antibiotics and antimycotics for prevention of bacterial and fungal growth, and the sample was placed immediately onto ice. In total, 237, 23 and 48 samples were available for analysis from Maale Hahamisha, Mevo Horon, and Mikve Israel, respectively. From this total captured population, we chose a subset of 29 samples to send for Next Generation Sequencing. In preparation for sequencing, samples were spotted onto AniCards (https://www.anicon.eu/card) for RNA preservation. Next Generation Sequencing, while much pricier than classical species-specific tests (e.g. swabbing and growing campylobacter colonies), represents a major leap forward in analyses of microbial composition, and allow us to link these observed loads across many species with relevant ecological predictors. Validating NGS result with *Campylobacter* and *Salmonella* colony growth shows a strong agreement between methods, with both approaches revealing relatively high prevalence of the former and negligible prevalence of the latter (Crafton et al, in preparation). All capture and sampling procedures were authorized by research permit 2019/ 04-19-059 provided by the Tel Aviv University ethics committee.

### Next-Generation Sequencing

To present a more in-depth comparison between movement and the microbiome, a total of 29 samples (Maale Hahamisha: 19, Mevo Horon: 5, Mikve Israel: 5) were analyzed using Next-Generation Sequencing by AniCon Labor GmbH in Germany (https://www.anicon.eu/). AniCon is a private company that discloses their full protocols, thus here only we briefly overview these analytical protocols of total RNA sequencing. Readers are also referred to the Supplementary Materials for additional details and to previous studies applying a similar approach with highly-specialized protocols of NGS swab preparations (Ayala et al. 2016, Dimitrov et al. 2017, Ferreira et al. 2019).In brief, the AniCards were first applied with the appropriate volume of sample media, properly dried, individually packaged in zip-lock bags with desiccants, and later sent for NGS.

In the lab, DNA was digested from the AniCard samples, and total RNA was extracted. After reverse transcription and inclusion of barcodes and sequencing adapters, the samples were analyzed for library preparation in a total RNA seq workflow. Random primers were used for RT after DNase digestion on a MiSeq machine in paired read mode, with a read length of 150 bp. This RNA sequencing approach was chosen because it outperforms 16S amplicon sequencing and shotgun metagenomics, by facilitating detection of viruses and bacterial sequences outside of the 16S locus (and therefore in most cases has higher discriminatory power than other sequencing methods). These methodologies represent cutting edge molecular and bio-informatics techniques capable of identifying microbiome composition at a higher resolution compared to traditional methodologies (Hempel et al. 2022). Accordingly, these protocols require more resources, thus limiting the sample size.

### Bioinformatics Analysis

**Taxonomic profiling of microbial agents** (pathogens and others): organisms in the data obtained from the MiSeq sequencing of the KAPA libraries were profiled at BASE2BIO LLC (Madison, WI, USA) by untargeted metagenomics discovery (avian) workflow. The pre-processed reads were searched against a custom database composed of known microbial sequences, host sequences and common contaminants. Data were filtered to reduce false positive hits based on cutoffs determined by bootstrapped simulations. Final classifications were filtered based on unique k-mer count (number of k-mers that can be assigned uniquely to that taxon or below) and adjusted read count or a-score, to which an adjustment factor was applied accounting for highly abundant species in a closely related clade. Each of the assembled contigs were annotated with the lowest common ancestor, which can be assigned to the sequence based on BLAST searches (GenBank). This produced datasets with two different stringency levels: “relaxed” and “strict”. For this particular study, we chose to analyze only reports of bacterial agents (i.e. excluding viral and fungal data from subsequent analysis) produced with “strict” cutoffs for k-mer count of 200 in order to limit the potential of relatively high numbers of false positives. The endpoint of this cutting-edge array of methods and analytical bio-informatics methods is the ability to identify the list of pathogens (and other agents) associated with each individual pigeon. Full details of bioinformatics protocols can be found in Supplementary Materials.

### Data Analysis of Movement

First, to appropriately represent individual’s movements we filtered GPS data by removing erroneous localizations, and excluding fixes with the following criteria: HDOP>2.1, Altitude > 2000, Number of Satellites <5. Additionally, we excluded tracking days with less than 20 points per day and individuals with less than 14 days of tracking, leaving a total of 33 pigeons for the analysis of movement data. For ensuring comparable movement indices we unified sampling rates across all individuals by sub-sampling data to fixes taken every hour.

To discover among-site and among-individual differences in movement or in the associated bacterial abundance, we extracted a few relevant movement indices that represent the spatial extent of the individual activity (namely max daily displacement and daily travel distance), and potential exposure to different locations (number of stops at unique locations). First, we calculated ‘Max Daily Displacement’ as the straight-line distance between the pigeon’s first location of the day (typically at the roost at dawn) and most distant point of the day. ‘Mean Daily Travel Distance’ is the sum of the straight-line segments between all consecutive GPS points of the day. To determine exploration level we quantified the daily number of spatially different stops each pigeon made. Each localization was considered a unique “stop” if it stayed longer than 30 minutes in the same location, and if this was at least 200-meter away from the last stop. A Kruskal-Wallis test was performed to determine if any of our movement metrics (average across all days for each individual) were significantly different between capture sites. In addition, we also used linear mixed models with pigeon ID as a random factor to determine the combined influence of capture location, sex, and weight. Since the latter two predictors had only very weak influence on movement in our dataset, these results are presented in the appendix.

### Microbiomes’ Alpha Diversity and its association with movement

To identify core bacteria in our NGS sample population at all three sites (n=29), we filtered the top 20 taxa present in at least 90% of the pigeons and visualized them in a Venn diagram, grouped by site, using the microbiome (Lahti et al, 2017) and Euler packages in R. In the supplementary materials, we also provide a heat map of this subset (**Figure S2**).

We investigated how pigeons and sites differed in their microbial diversity. We performed several exploratory analyses and differential diversity analyses of pigeon microbial profiles in R using the Phyloseq package (McMurdie and Holmes 2013) with a log base 10 transformation to normalize the data. There are 3 available indices of richness (Observed, Chao1, ACE) and 4 of diversity (and Evenness: Shannon, Simpson, Inverse Simpson, Fisher’s alpha). Our results of all indices largely agreed with each other (**Figure S3)**; for simplicity, we present here results for one index of each currency: Chao1 for richness and Fisher’s alpha for diversity. The Chao1 metric for richness estimates the number of different bacterial genera in a sample and is particularly useful for data sets skewed toward the low-abundance classes, as is likely to be the case with microbes (Chao et al. 2006).

Fisher’s *a* index is recommended as a reliable measure due to its ability to compensate for potentially incomplete sampling (Hayek 1997) ; it is relatively unaffected by variation in sample size and aims to measure diversity by accounting for “evenness” or homogeneity. For instance, communities dominated by one species will exhibit low evenness, as opposed to communities where most species are relatively well represented (high evenness).

Focusing on the association between movement and bacterial abundance (while accounting for differences among sites, and individual traits) is possible for the subset of pigeons with both sufficient GPS movement data and NGS data. This subset included 20 individuals, for this particular analysis. Here, we have used general linear models (GLM) with the package LME4 (Bates et al. 2015) to test whether capture location, sex, body weight (a possible indicator of condition), and movement indices predict differences in Chao1 and Fisher diversity among pigeons. To assess the accuracy and performance of multiple models, the Akaike information criterion (AIC) was used to rank competing models, and the model with the lowest AIC value was reported. All second-best models which had considerably lower scores were included in the supplementary materials.

### Microbiomes’ Beta Diversity and its association with movement

Beyond the effect on alpha diversity, we also explored the effects of site and individual sex and movement indices (Daily Max Displacement, Daily Mean Travel Distance, Average Number of stops) on beta diversity. Here we implemented a non-metric multidimensional scaling (NMDS) and redundancy analysis (RDA) using the *vegan* package. Broadly speaking, redundancy analyses combine methods of linear regression analysis with methods of principal components analysis to predict how specific variables affect a response (e.g. site and movement indices of each individual). Because most linear methods (including RDA) are sensitive to the distribution of the data - and specifically to the many zeros that characterize microbiome count data - The Hellinger transformation of relative abundance is recommended as a preliminary step (Legendre and Gallagher 2001). The significance of separation in RDA analysis was estimated with a permutation test using 5,000 permutations. Distance-based comparisons among individuals were made using the Bray-Curtis similarity coefficient calculated between each and every sample pair and visualized with nMDS and RDA. ANOVA was used to test for the significance of movement indices on beta diversity using an alpha level of 5%.

## Results

### Pigeon movement

Out of the 308 color-ringed individuals, 102 were re-sighted during our routine fieldwork. A total of 33 pigeons with adequate GPS data (out of 44) accumulating to 8635 days (214±193 range 14-713 days per individual) of tracking were analyzed for Daily Mean Travel Distance, Max Daily Displacement, and Average Number of Stops (**Table 1**). Mean Daily Travel distance was significantly different between sites (*F*=4.0, *df*=2, *P*=0.002). Specifically, Mikve Israel (n=8) differs from Mevo Horon (n=6, *P*=0.004), and Maale Hahamisha (n=19, *P*=0.004; **Figure 2**, **Table 1**). Mevo Horon and Maale Hahamisha did not differ (*P*=0.465). We found that Max Daily Displacement also differed between sites (*F*=6.2, *df*= 2, *P* = 0.009). Similarly to Mean Daily Travel Distance, Mikve Israel was significantly different from Maale Hahamisha (*P*= 0.03), and from Mevo Horon (*P*=0.01), but Maale Hahamisha and Mevo Horon did not differ (P=0.272; **Figure 2**). Pigeons captured at Mevo Horon averaged the largest Max Displacement (3.29 km) and Daily Travel Distance (2.50 km) compared to the other two sites (**Figure S4**). In contrast to the pronounced differences between sites in movement ranges, the average number of stops (our index of exploration) differed only slightly between them (*F*=5.9, *df*= 2, *P* = 0.05) with a large variation among individuals within sites. Parameter estimates for the top model used for GLMM describing movement indices can be found in the appendix **(Table S2**). Additionally, sex and weight did not significantly affect movement in our dataset (Appendix **Table S3**).

**Figure 2.**
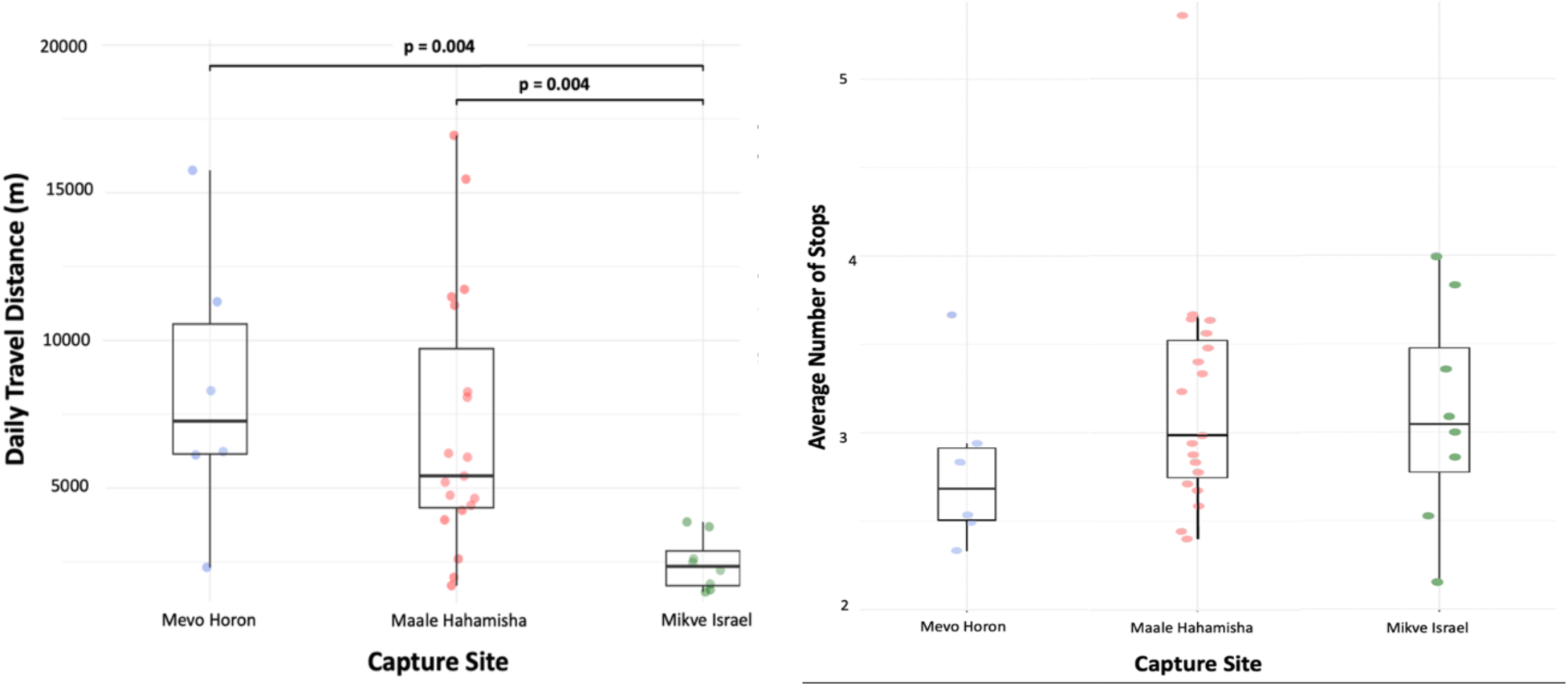
Pigeon movement within the three study sites. (**A**) *Mean Daily Travel distance; and (**B**) Average Daily Stops* among sites ordered by urbanization level: Mevo Horon as our most rural site, Maale Hahamisha semi-rural, Mikve Israel highly urban. Bars indicate significant differences between specific sites. While pigeons in Mikve Israel traveled shorter distances compared to other two sites, variation in daily Number of Stops differed only slightly between sites.

**Table 1.** Tracking data and basic movement characteristics in the three study sites (Mean ± sd).

| Capture Site | *n*<br>pigeons | Tracking<br>duration<br>(days) | Mean Daily<br>Travel<br>Distance<br>(m) | Max Daily<br>Displacement<br>(m) | Exploration<br>(average<br>number of<br>Stops) |
| --- | --- | --- | --- | --- | --- |
| **Mevo Horon**<br>**(rural)** | 6 | 283.8+65.8 | 8,339+1,910 | 3,672+934 | 2.80+0.19 |
| **Maale Hahamisha**<br>**(semi-urban)** | 19 | 300.7+46.3 | 7,061+1,008 | 2,516+521 | 3.18+0.15 |
| **Mikve Israel**<br>**(urban)** | 8 | 152.3+25.6 | 2,449+322 | 966+235 | 3.10+0.21 |
| **Total** | 33 | 245.6+33.0 | 5,949+764 | 2,384+375 | 3.02+0.16 |

### Pathogen presence in pigeon microbiome

To compare the core taxa shared between the three study sites, we identified the top 20 bacterial genus found in 90% of samples within each site, with a relative abundance higher than 0.1% (**Figure 4.A**; see **Table S5** for a full list of identified genera). The majority of these most-prevalent genera (found at all sites) such as Clostridium, Escherichia, Campylobacter, and Streptococcus contain many species of disease-causing pathogens. A number of these species were identified in our NGS reports, and here we include those which have known cases of associated illnesses in humans, cattle, and poultry (**Figure 4.B**), and thus demonstrate pigeons’ potential to carry relevant pathogens at our population.

**Figure 3.**
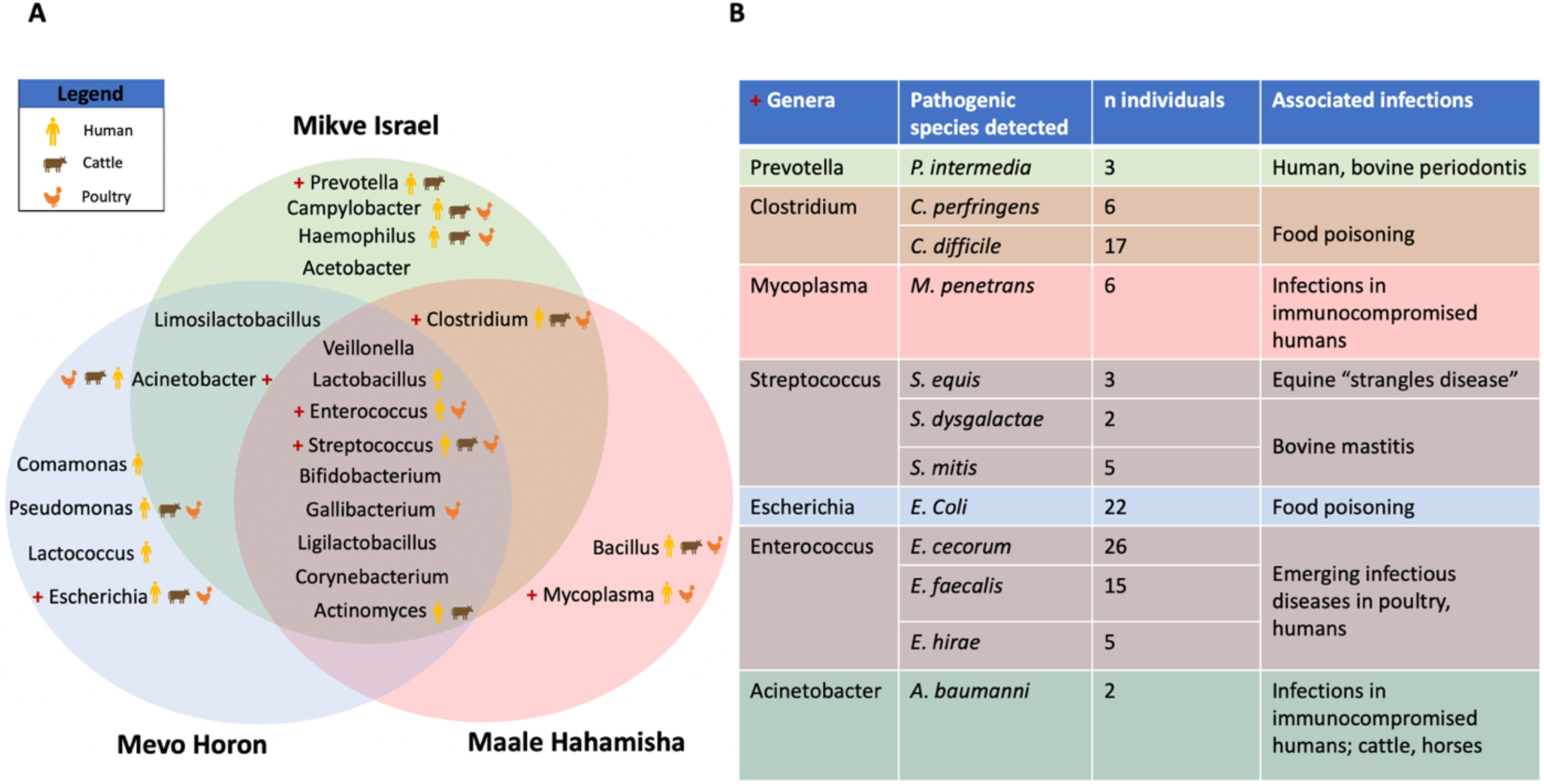
**A)** Top 20 taxa at the genus level (filtered by highest abundance and identified for at least 90% of samples at each site), by their occurrence in the different study sites. Genera with known pathogenic species (one or more) within them to either Humans, Cattle or Poultry are labeled. **B)** List of individual species with known pathogenic potential identified from NGS reports within Top 20 genera.

**Figure 4.**
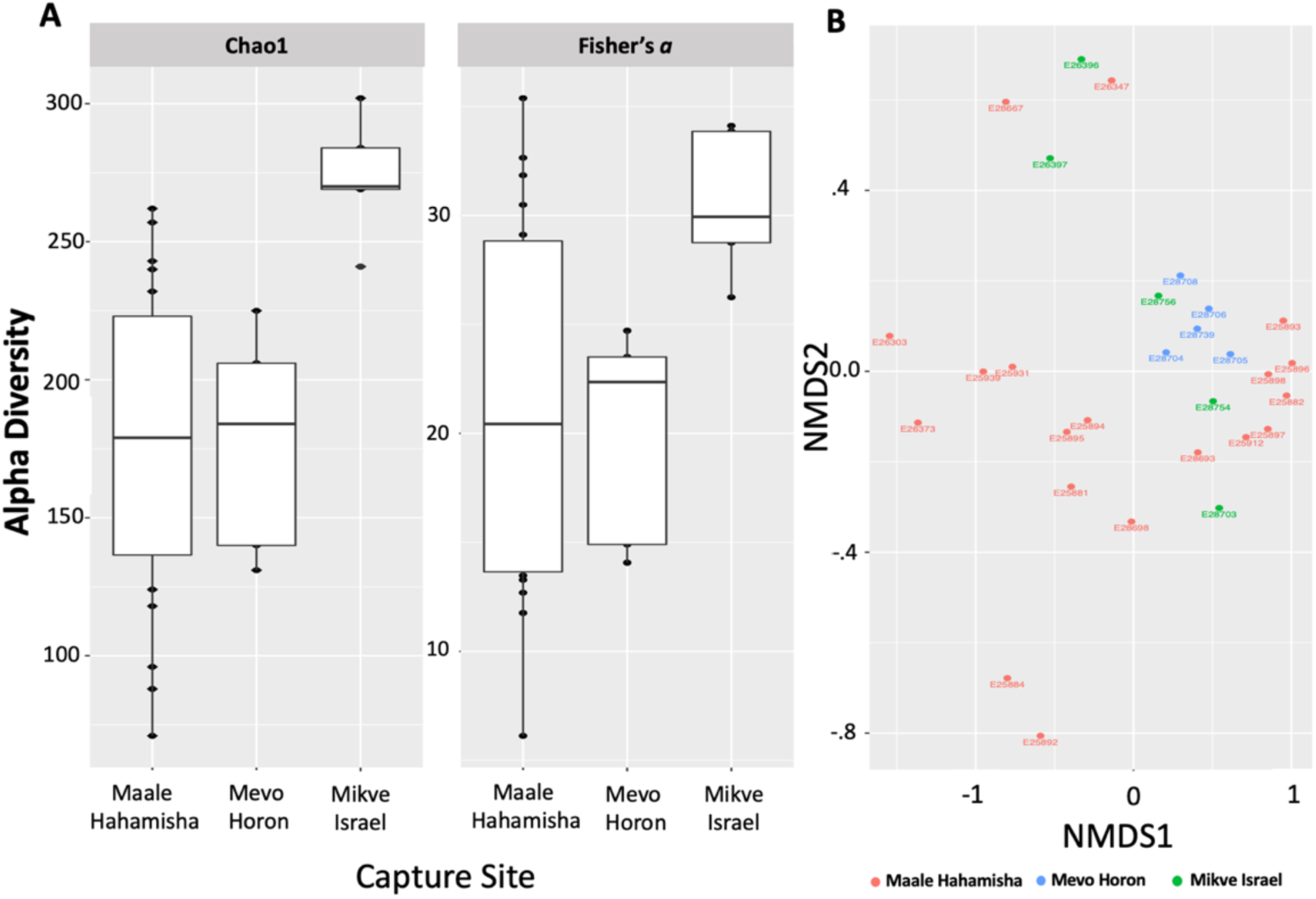
(A) Microbiome diversity across the three study sites. (**A**) The comparison of microbiome alpha diversity between sites, including genera richness (represented by Chao1) and evenness (represented by Fisher’s index). The microbial community show that pigeons in Mikve Israel (the most urban site) had higher richness and evenness than the two rural sites at Maale Hahamisha and Mevo Horon. **(B)** Beta –diversity, non-metric multidimensional scaling (nMDS) ordination of pigeon microbiome communities within each site. Circles represent individuals, and distances between individuals represent similarities between samples (closer individuals are more similar than distant individuals).

### Effect of site on Alpha and Beta microbiome diversity

First we tested alpha diversity between capture locations. Fisher’s alpha diversity index was marginally significant (Kruskal-Wallis *P* = 0.05). The Chao1 index for richness showed a significant variation among capture locations (Kruskal-Wallis *P* = 0.004), with pigeons from Mikve Israel showing significantly higher richness than pigeons from both Mevo Horon (posthoc Dunn’s test, *P* = 0.01) and Maale Hahamisha (posthoc Dunn’s test, *P* = 0.004) (**Figure 4**). This is congruent with other richness metrics (Observed, ACE) suggesting that pigeons in Mikve Israel (highly urban) contain a higher number of lower abundance species as opposed to pigeons from Mevo Horon (highly rural) and Maale Hahamisha (semi rural) that did not substantially differ from each other. Chao1 index provides an estimate of the expected number of species in a habitat.

### Effects of movement and capture site on microbiome alpha diversity

After filtering for pigeons with both NGS data and sufficient GPS movement data we were left with a total of 20 individuals for an analysis of the relationship between movement and bacterial abundance. Interestingly, when modeling jointly the effect of site and movement on alpha diversity/richness we found that the average number of stops (*F*=6.02, *df*=2, *P*=0.02) and capture site (*F*=4.02, df=2, *P*=.02), but not their interaction (*F*=1.9, df=2,*P*=0.47) were significantly correlated to alpha diversity (**Table 2**). Beyond the clear differences among sites (∼10 more genera in Mikve Israel compared to the other two), within each site pigeons that regularly made more stops per day (to different unique locations) show higher alpha diversity. This finding implies that the higher environmental exposure by individuals that visit more diverse locations indeed results in a more diverse microbiome in both richness and diversity.

**Table 2.** Effects of site and movement patterns on alpha diversity and species richness indices. Generalized Linear Models for Chao1 and Fisher’s alpha showing that feral pigeons in central Israel varied in their microbiome alpha diversity. Effects of daily travel distance, max daily displacement, and the interaction between number of stops × capture site were performed in each model, but were not found significant and therefore not included in the table. Significant *P* values are in boldface.

| Diversity index | Factor | Estimate | Std. error | p. value |
| --- | --- | --- | --- | --- |
| Chao 1 | ***Movement*** |  |  |  |
|  | Number of stops | 37.75 | 13.44 | **0.012** |
|  | ***Capture Site*** |  |  |  |
|  | Mevo Horon | 13.46 | 24.94 | 0.596 |
|  | Reference (Maale Hahamisha) | 60.15 | 52.38 | 0.261 |
|  | Mikve Israel | 102.12 | 23.25 | **0.0004** |
| Fisher's alpha | ***Movement*** |  |  |  |
|  | Number of Stops | 5.59 | 1.90 | **0.023** |
|  | ***Capture Site*** |  |  |  |
|  | Mevo Horon | 0.278 | 3.28 | 0.938 |
|  | Reference (Maale Hahamisha) | 2.91 | 7.89 | 0.718 |
|  | Mikve Israel | 9.98 | 3.52 | **0.007** |

The lack of interaction implies that this positive association was similar at all three sites (**Figure 5**). Mean Daily Travel distance was significantly different between sites, (Mikve Israel vs. Mevo Horon), but was not correlated with variation in alpha diversity among pigeons within each site (*F* = 0.18, df=2, *P*=0.71) nor did capture site and Mean Daily Travel Distance interact in their effect on diversity (*F* =0.13, *P* = 0.84). Similarly, Max Daily Displacement did not affect alpha diversity beyond the site effect. Sex and body weight did not have a significant effect on movement or microbiome diversity and are therefore not presented in the main text but can be found in the supplementary materials section (**Table S4**).

**Figure 5.**
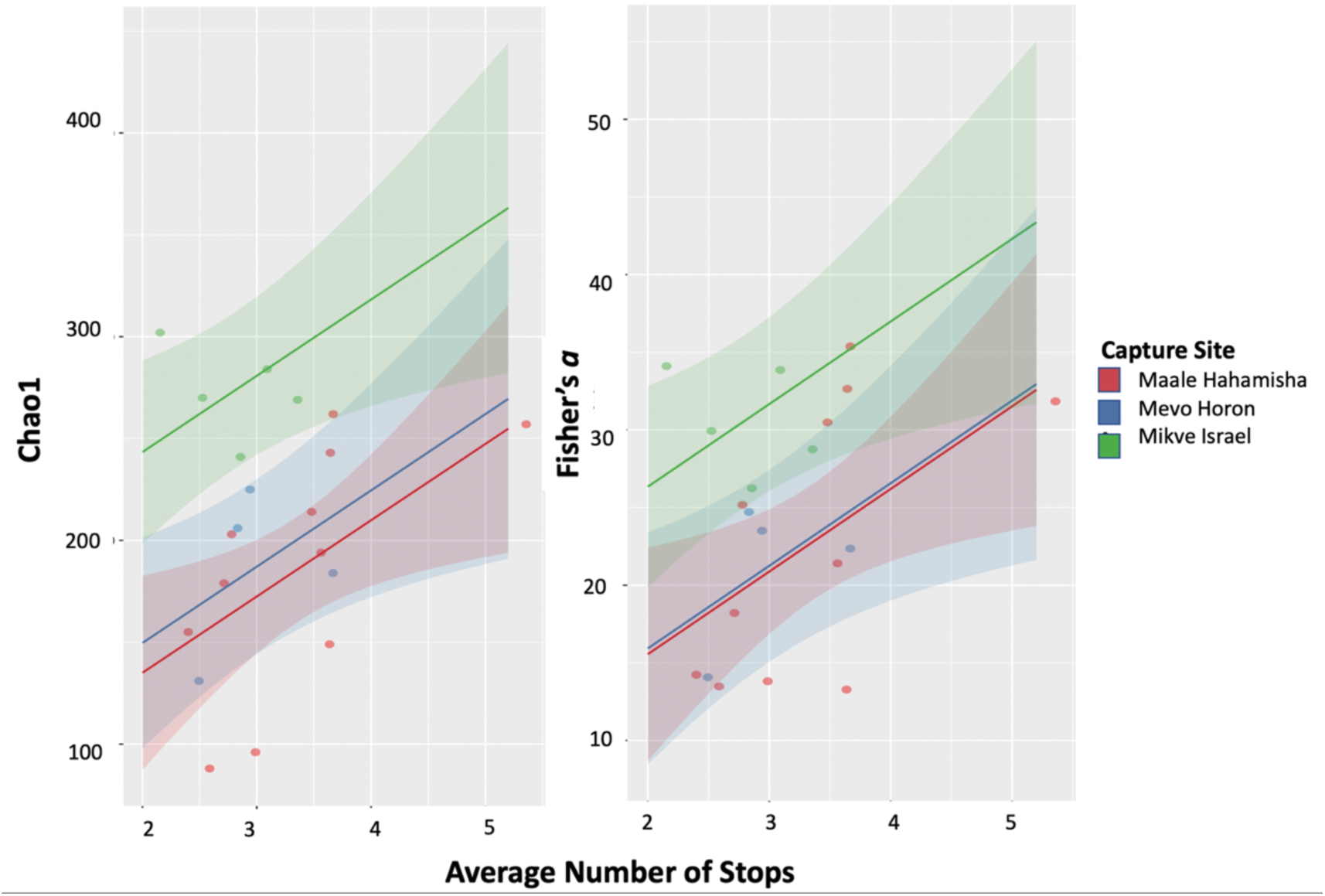
The effect of pigeon movement on microbiome diversity. Results of general linear modeling (GLMM), with the response variable of Chao1 for diversity and Fisher’s alpha for evenness, showing higher bacteria evenness associated with more daily stops. Shaded areas represent the smoothed 95% confidence interval. Results are robust to the removal of the outlier (see Figure S4).

### Effects of movement and site on microbiome beta diversity

Focusing on Beta diversity, the RDAs models (**Figure S6**) suggested that individuals from different capture locations tended to be colonized by different bacterial taxa (*F* = 1.357, *R^2^* = 0.102, *P* = 0.043). In addition, average number of stops had a significant effect on community composition (*P*=0.007), while sex and Mean Daily Travel Distance were not significant (P=0.758, *P*=0.400). These results agree with the results mentioned above for alpha diversity.

## Discussion

In this study, we explored the role of feral pigeons as potential pathogen vectors at dairy farms and the association between movement and microbiome. First, GPS tracking showed that pigeons on the urban farm (Mikve Israel) were largely less mobile and more local than those in our two rural sites and that individuals (within sites) differed in their movement patterns (**Figure 3**). Second, NGS results were consistent with previous literature, showing that the pigeons in our population have a high prevalence of pathogens relevant to human, cattle, and poultry. The top 20 most abundant taxa include Escherichia, Campylobacter, Clostridium and Streptococcus, and several other genera that contain important pathogenic species of bacteria which were identified in our NGS reports (**Figure 4**). Third, pigeons’ microbiome differed among sites with higher alpha diversity and richness found in the urban site (**Figure 5**). Further, pigeons’ movement patterns (within each site), but not their sex or weight affected their microbiome diversity –explorer individuals who stop at more diverse places show a corresponding increase in microbiome diversity beyond the effect of sites (**Figure S6**). To the best of our knowledge, these links between spatial behavior studied *in situ* and bacterial diversity in wildlife were only rarely investigated (e.g. in Fuirst et al. 2018), and are especially missing from studies in the context of agricultural One-Health approach. The finding that pigeons’ potentially transmit pathogens among dairy farms, human settlements (where they almost exclusively roost), and wildlife (where they often forage) highlights the potential importance of their relationship with surrounding habitats. The association between exploratory behavior and epidemiology in our analyses of movement and microbiome diversity directly offers an example of the link pigeons create between human and animal populations, (and their pathogens), in urban and agricultural sites. Below we discuss differences in pigeon movement between sites, then how each site differs in microbial diversity, the effect of pigeon movement on diversity, and finally the broader implications of our findings.

### Variation in movement patterns and shorter distances in urban pigeons

First, we studied whether there were differences in movement between individuals and sites, addressing the spatial extent of pigeon transmission potential and its variation with respect to urbanization. Pigeons captured at each site were active in its vicinity and crossing among sites was very rare. Mevo Horon and Maale Hahamisha are more rural than Mikve Israel, with Mevo Horon being slightly more isolated. Accordingly, pigeons daily travel distances and displacement were longer at these two sites compared to Mikve Israel, likely reflecting different resource distributions among sites. All pigeons roosted in buildings within human settlements and routinely visited the respective dairy farms and similar anthropogenic food sources (e.g. parks, poultry farms, and commercial areas). Yet, rural pigeons spent summer months foraging for grains at harvested fields, while the urban ones rarely did so (and only in a few nearby fields). Instead, they foraged year-round on human waste. Thus, pigeons foraging at Mevo Horon must travel further from their urban roosting locations when searching for surrounding food, offering a likely explanation for higher displacement when compared to other sites. These patterns agree with general findings that urban animals move less thanks to the availability of stable and rich food sources (Tucker et al. 2018) .

Identifying heterogeneities in host space use – here among sites as discussed above - offers important insights toward understanding pathogen transmission dynamics (Dougherty et al. 2017). Pigeons can spread pathogens over a range of agricultural and urban habitats, affecting livestock, wildlife, and humans. For instance, non-resident pigeons in cattle farms in Colorado are responsible for pathogen transmission among farms, and the resident ones for amplification within farm (Carlson et al. 2011). The significantly larger travel distance and displacement of the pigeons foraging at Mevo Horon and Maale Hahamisha, as well as the composition of habitats used, might suggest a more widespread range of transmission potential when compared to pigeons foraging at Mikve Israel (**Figure 3**). The pigeons at Mikve Israel, on the other hand, might pose a greater risk for zoonotic diseases due to their high reliance on urban centers and dairy farms and the high rate of alternation between the two, sometimes located within hundreds of meters. Further research is needed to directly address this ‘urbanization-dependent transmission potential’, but nevertheless this hypothesis is supported by a growing consensus that variation in host home range has the ability to influence disease spread (McClure et al. 2020) .

### Pigeons from different sites differ in their microbiome diversity

Our results show that pigeons captured at Mikve Israel had significantly higher microbial alpha diversity compared to both Mevo Horon and Maale Hahamisha (Table 2, **Figure 5**). Each pigeon in Mikve Israel had around 160 genera of bacteria (Chao1) and diversity of ∼13 (*Fisher’s a*) compared to those in the latter two sites with a similar diversity of around 70 and 3, for the two indices respectively. Admittedly, working with merely three sites that may differ in various aspects (e.g. local ground elevation, resource composition, local density, and others) does not permit certain identification of the main cause for this result. Nevertheless, the urbanization gradient is an apparently obvious possible explanation for this finding. Indeed a substantial positive effect of urbanization on microbial diversity and the microbial communities of wildlife was attributed to acquiring parasites, natural infections, and other physiological stressors from urban habitat (Rouffaer et al. 2017).

However, the literature is inconsistent with respect to the effect of urbanization on microbial diversity, and the opposite (negative) trend was also found in synanthropic birds (Jatzlauk et al. 2017). For instance, Fuirst et al (2018) studied the effects of urbanization on foraging and microbiota in three gull colonies (*Larus argentatus*) and found that bacterial diversity was highest in the colony that had the *lowest* urban exposure. Such disagreement can either reflect a genuine variation between systems in their response patterns to urbanization or methodological differences. Indices used for urbanization and diversity, the sampling methods, and the spatiotemporal scale of investigation can all act as confounding factors.

### Individual host movement is correlated with the alpha diversity in their microbiome

Beyond the variation among sites, more exploratory pigeons from all sites that averaged more stops in unique locations (i.e., excluding repeated stops within a day at the same location) had higher richness and diversity than individuals that typically spent their day foraging in fewer places (e.g. a dairy farm; **Figure 5**). These results agree with previous findings indicating the association of microbiota with foraging patterns in other birds (e.g. Corl et al 2019, Pekarsky et al. 2021). In agreement with our result, also the gulls in the above-mentioned study by Fuirst et al (2018) had the maximal bacterial diversity in the colony that used a wider variety of foraging sites (i.e. lower foraging site fidelity).

Nevertheless, it is noteworthy that this colony also had the *lowest* urban exposure among the three colonies included in their study. This coupling between urbanization and foraging behavior impairs the ability to dissect between the independent effects (if any) of foraging and urbanization in their system. In our case, the number of stops was similarly spread in all three sites (**Figure 3**), allowing us to dissect between the effect of (site) urban level and (individual) foraging behavior.

Why should individual behavior be correlated to their microbiome diversity? There are at least two broad explanations for this finding. First, it may indicate that individuals that utilize an increased diversity of spaces encounter a higher diversity of bacteria on route (i.e. sample their region more thoroughly). Fuirst et al (2018) for instance, suggested this explanation – the diverse foraging locations resulted in a more diverse microbial community. An alternative mechanism suggests that the microbiome composition itself affects pigeons’ behavior (and not vice versa), through sickness or host manipulation done by one or a few of the agents within the microbiome (Poulin and Maure 2015). It is possible that individuals with higher microbiota evenness and diversity may need to visit more locations in order to meet their nutritional requirements. For instance, a negative relationship was found between gut diversity and home range size of white-tailed deer (Webb et al. 2010). The authors suggested that deer with higher gut microbiota evenness are able to use more diet sources overall (i.e. the gut bacteria allowed for eating different plant species) and thus can meet their nutritional requirements within a smaller area relative to individuals with less diverse gut microbiomes (Webb et al. 2010). Importantly, these two explanations can be non-mutually exclusive or vary in their relative contributions across different systems and ongoing studies will further illuminate these pathways (Couch and Epps 2022).

### Concluding remarks

The combined effects of high mobility (daily ranges of a few kilometers), high densities (hundreds of individuals in each site), high proximity to humans and cattle (at roost and foraging sites, respectively), as well as opportunistic visits to open areas, point to the very strong potential of feral pigeons to directly transfer pathogens between livestock, humans, and wildlife. Understanding host microbial diversity allows critical insights and identification of important viral and bacterial pathogens that may have potential to cause devastating human pandemics; for example three global plagues caused by *Yersinia pestis*, and a Cholera pandemic caused by *Vibrio cholerae* (Sacchi et al. 2002).

Other synanthropes are likely to play a similar role in a variety of anthropogenic systems. While the scope of this study is human-centered with the concern of zoonotic disease, these patterns should be considered also for conservation and spillover effects since pigeons are similarly likely to transmit pathogens in the opposite direction, risking wildlife populations (e.g. while foraging at mixed flocks in open fields). The massive outbreak of Avian Flu in 2022 (H5N1) likely originating from poultry farms (Center for Disease Control, 2022) and infecting many endangered species, is a sad reminder of this scenario.

In addition to the obvious agreement with the main premise of the One Health approach (e.g. Hassell et al. 2017), the remarkable differences among and within sites (in both local diversity and movement patterns), emphasize the need to adapt response strategies tailored to local conditions and investigate focal systems towards a mechanistic understanding of these processes. For instance, preventive pigeon culling, if applied in rural areas, should be done in concert across much larger areas than their more urban counterparts. Similarly, within sites, more exploratory pigeons should be targeted when possible (e.g. by trapping at larger distances from the center of activity or prioritizing individuals on the move). Amending practical culling protocols with such insights regarding wildlife movement can indicate how movement ecology studies can facilitate applied disease management (Dougherty et al. 2017, Spiegel et al. 2017).

Our study is simplistic in several aspects; first sample size limitations imposed by the high costs of both tracking devices and NGS processing prevent investigating the question across a larger set of sites (e.g. better representing a gradient of urbanization) and accounting for additional factors such environmental conditions (e.g. seasonality) and demographics (e.g. age-related differences, density, social contexts) and others. Similarly, the study suffers from an unbalanced design with better representation of Maale Hahamisha, while poorer coverage of the other two sites. This bias reflects both historical reasons in the study development as well as methodological challenges with the ability and permits to work in the latter two sites. While we acknowledge this as an important consideration to the study, we do not believe it invalidates our results, and reported effect size and certainty levels reflect these variable sample sizes. Second, while our exploratory analyses allow us to determine the co-variation between simple indices of movement and bacterial diversity (higher in more exploratory pigeons), they are short from targeting specific mechanisms or identifying how specific pathogens are affected by differences among sites and behaviors. Future studies can build on these general patterns and proceed beyond our observational approach to identify the causality of this positive correlation.

Experiments like inoculation, translocations, movement restrictions, and other *in situ* manipulation can reveal the relative contributions of higher exposure (i.e. movement drives pathogens prevalence) vs. pathogenic impact on movement (e.g., different needs or host manipulation). Assimilation of tools and concepts from movement ecology into studies of disease ecology is a promising direction toward this end (Dougherty et al. 2017; Ezenwa et al. 2022).

## Acknowledgments

We acknowledge financial support by our funders Grant 891-0232-21 from the Israel Dairy Board Research, and grant ISF396/20 from the Israeli Science Foundation.

We gratefully acknowledge members and workers of Maale Hahamisha, Mevo Horon, and Mikve Israel for allowing us access to the study sites and providing assistance in the field. We thank members of the Spiegel lab (Nili Anglister, Eitam Arnon, Hilla Ziv, Yohay Wasserlauf, Tovale Solomon) for their help with data analysis and fieldwork. A special thanks to Henning Bischoff of AniCon Labor GmbH, Jeremy Volkening, and Claudio Afonso of Base2Bio (Oshkosh, WI, USA) for support with NGS data and the manuscript.

## Author contributions

MC, SC, and OS initiated the study; OS, AL, and SC secured the funds; MC, SC, and OS did the fieldwork, MC and AL did labwork; MC and OS analyzed the data with inputs from SC, AL, LR, CA; MC and OS wrote the manuscript with a substantial contribution from all other authors who approved the final version.

