## Supplementary Materials for "Association between movement patterns, microbiome diversity, and potential pathogen presence in free-ranging feral pigeons foraging in dairy farms"

### Methods

### Urbanization level analysis at each site

Our sites (Mevo Horon, Maale Hahamisha and Mikve Israel) lie on an obvious gradient of increasing urbanization. To quantify this intuitive notion, we used Google Earth areas photos to create polygons for each urban area within a 2km diameter surrounding each site (a scale relevant to pigeon's movement, see results). All area that did not fall inside urban polygons (namely, non-built areas) was considered rural. These polygons were analyzed in R to calculate what percent of the area surrounding the site is considered urban, or rural. Results agree with the qualitative estimation, with zero percent of the land within 2km surrounding Mevo Horon considered urbanized, therefore making it our most rural site. Maale Hahamisha, had merely 13.27% of the surroundings being urban. Finally, Mikve Israel had 41.97% of its surroundings considered urban, thus making it the most urbanized site.

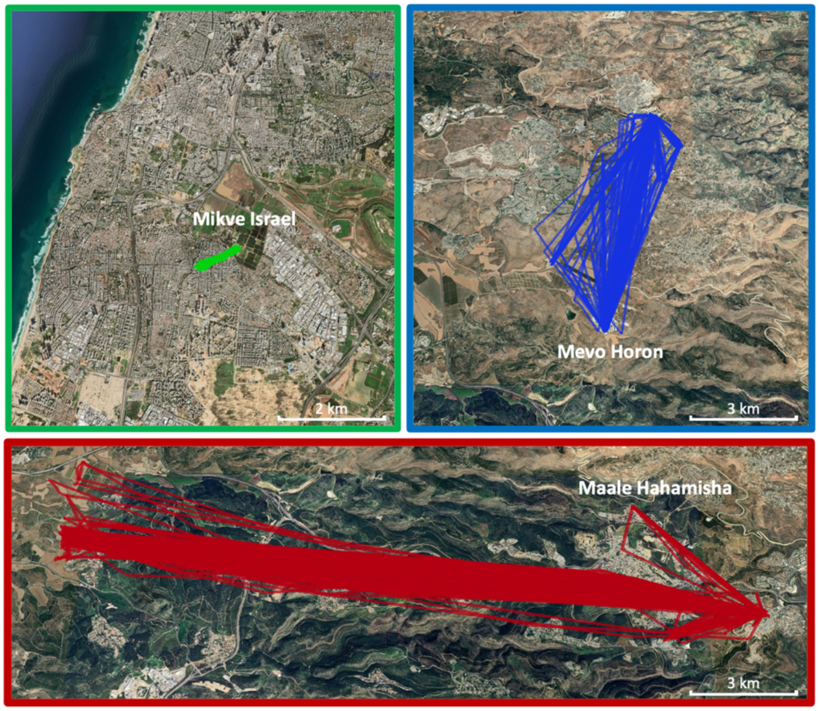


**Figure S1. Travel distance variations between sites.** Randomly chosen examples of a typical individual from the three sites. Pigeon E28743 (an adult male) captured at Mikve Israel shows a travel distance of around ~1km, while pigeon E28759 (an adult male) at Mevo Horon a typical daily distance of 6 km, and the one at Maale Hahamisha (pigeon E28688, an adult male) travels around ~15km.

### NGS

Next generation sequencing (NGS) was performed by AniCon Labor GmbH (https://www.anicon.eu/) using total RNA extracted from AniCards spotted with field specimens. While these are not the exact methods of AniCon, here we describe similiar swab preparations and the highly specialized protocols of RNA Sequencing. More detailed descriptions can be found in previous literature (Ayala et al. 2016, Dimitrov et al. 2017, Ferreira et al. 2019).

**Swab preparation**: Oral and cloacal swabs are spotted following manufacturer instructions with a volume of media, properly dried, individually packaged in zip-lock bags and with desiccants. Three discs (3 mm) were randomly punched out from each of the spot using sterile disposable biopsy punches. Duplicates of 10 discs from the same card were pooled (in duplicates) in sterile Eppendorf tubes containing freshly-prepared 120 µl TE (10 mM Tris-HCl; 0.1 mM EDTA; pH 8.0), followed by incubation at room temperature for 30 min. Pathogens in the samples were inactivated by addition of 101-µl MagMax viral lysis buffer (Ambion, Life tech), and the materials transferred to a BSL-2 laboratory for further processing. Total RNAs were extracted from the samples using MagMAXTM-96 AI/ND Viral RNA Isolation Kit (Thermo Fisher Scientific, MA, USA). To optimize the recovery of extracted RNAs, eluates from one of each duplicate were used to elute RNAs extracted from respective duplicates. The total RNA was quantified by Qubit® 2.0 Fluorometer (Life Technologies).

**Preparation of NGS sequencing DNA libraries:** NGS libraries were prepared by KAPA RNA-Seq (KAPA Biosystems, Boston, USA) by heat-fragmentation (30°C for 3 min), followed by synthesis, random priming and marking of the first and second strand cDNAs. cDNA library fragments will be purified with 1.8X AMPure XP beads, A-tailed on the 3'-ends, followed by addition of adapters. The libraries were amplified using KAPA HiFi HotStart PCR and normalized by purification using AMPure XP bead. Then these libraries were eluted in 22.5 µL of 10 mM Tris-HCl buffer (pH 8.0).

**Illumina MiSeq sequencing of KAPA libraries:** The KAPA libraries were quantified by Qubit® and Agilent 2100 Bioanalyzer System (Agilent technologies Inc., Germany) and sequenced by MiSeq (300 cycles; Illumina, USA) using Escherichia phage phiX174 (Bacteriophage phi-X174) as an internal control (Dimitrov et al. 2017, He et al. 2018).

### Bioinformatics

Raw reads were trimmed using Trim Galore (https://github.com/FelixKrueger/TrimGalore) v06.7 to remove residual adapter sequences and low-quality 3' ends. After masking of low-complexity subsequences using BBTools (v38.96) (http://sourceforge.net/projects/bbmap), reads were assigned taxonomic classifications by k-mer search using KrakenUniq (v0.5.8) (Breitwieser et al. 2018) modified with local patches, against a hierarchical set of databases containing vector/contaminant sequences, host genome (*Columba livia,* Cliv_1.0), human genome (GRCh38.p13), and the $BASE_2BIO$ internal untargeted database of microbial reference sequences, consisting of a representative subset of NCBI microbial genomes. Classifications were further adjusted using a patched version of the 'krakenuniq-filter' script, adjusting assignments up the taxonomic tree until the k-mer specificity was 0.05 for viral taxa and 0.25 for all other taxa. Each identified taxon was then further verified by BLASTn (Altschul et al. 1990) search of a random subset of taxon-assigned reads against the full GenBank 'nt' database and subsequent lowest common ancestor (LCA) assignment by in-house tools. These results were used to assign a secondary concurrence score to each taxonomic ID, calculated as the fraction of subsampled reads assigned to that taxon via full BLASTn search which agreed with the k-mer classification-based assignment.

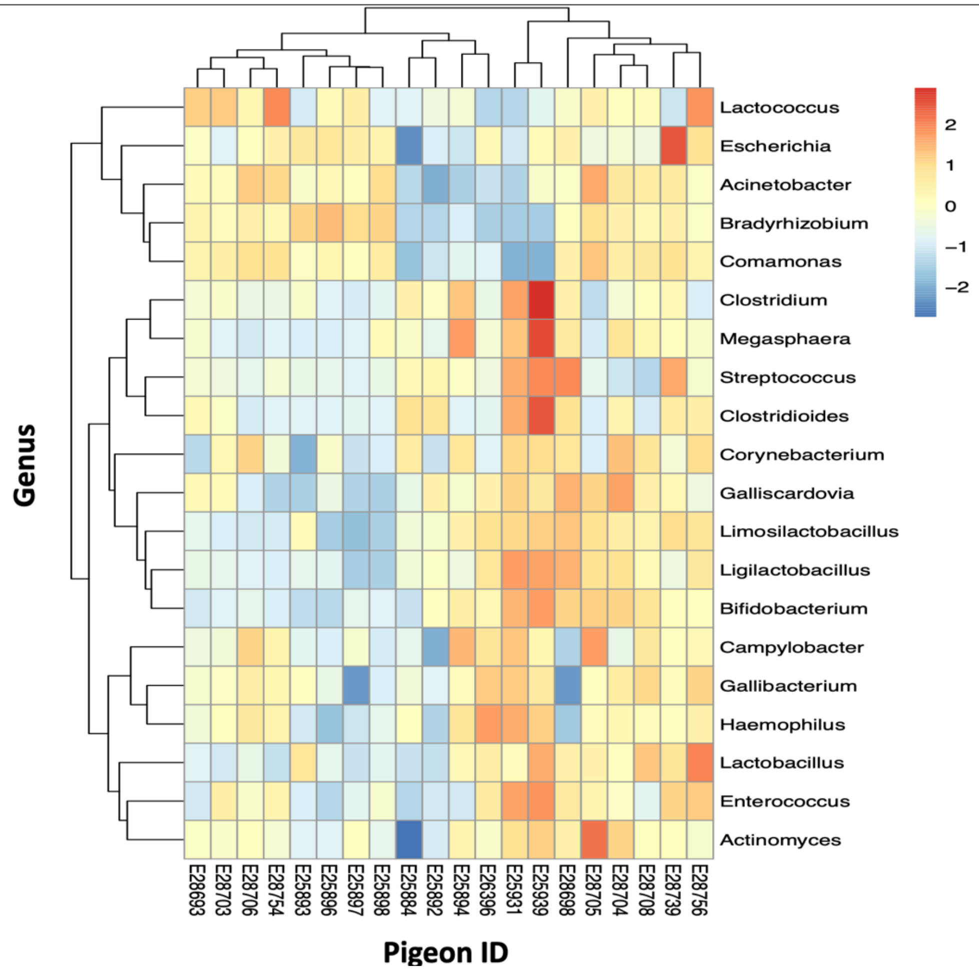


**Figure S2**. *Heatmap of top 20 taxa at the genus level, and relative abundance in all individuals.*

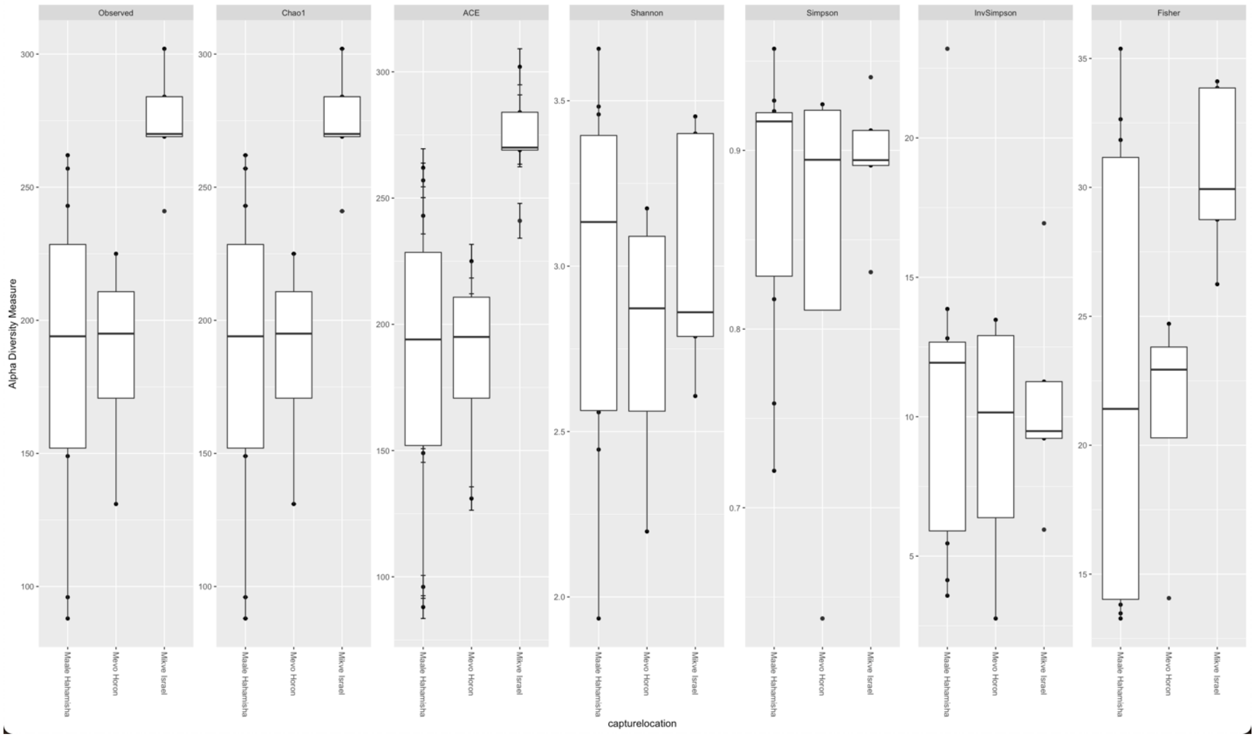


**Figure S3**. A graph including all diversity (Observed, Chao1, ACE, Shannon, Simpson, InvSimpson, Fisher) measures produced by the phyloseq package in R.

**Table S2. Parameters estimates for the top model used for GLMM describing movement indices**

| Movement index | Factor | Estimate | SE | p.value |
|---|---|---|---|---|
| Daily Max | ***Capture location*** | | | |
| Displacement | Mevo Horon | 1330.1 | 921.1 | 0.1579 |
| | Reference (Maale Hahamisha) | 2698.9 | 532.7 | 0.00014 |
| | Mikve Israel | -1434.6 | 835.8 | 0.0953 |
| Daily Mean | ***Capture location*** | | | |
| Travel Distance | Mevo Horon | 0.278 | 3.28 | 0.938 |
| | Reference (Maale Hahamisha) | 7143.2 | 861.1 | .0000000011 |
| | Mikve Israel | -4630.7 | 1624.0 | **0.007** |
| Average Number | ***Capture location*** | | | |
| of Stops | Mevo Horon | -0.503 | 0.316 | 0.895 |
| | Reference (Maale Hahamisha) | 3.540 | 0.158 | 0.805 |
| | Mikve Israel | -0.150 | 0.298 | 0.936 |

**Table S1. Linear mixed model comparison for the effects of capture-location, pigeon sex and weight on movement indices (max displacement, travel distance, and average number of stops).** AICc ranking of competing models show that the model with capture location as the sole fixed-effect was rank was top-ranked for all three indices. Sex and weight had minor influence on movement in our dataset.

| Movement index | Effect | AICc | Δ AICc | AICc Weight |
|---|---|---|---|---|
| Daily Max Displacement | Capture location | 528.25 | 0.00 | 0.64 |
| | Capture location + Sex | 530.88 | 2.63 | 0.82 |
| | Capture location + Weight | 531.18 | 2.93 | 0.97 |
| | Capture location + Sex + Weight | 534.12 | 5.87 | 1.00 |
| Daily Mean Travel Distance | Capture location | 574.72 | 0.00 | 0.69 |
| | Capture location + Sex | 576.75 | 2.03 | 0.93 |
| | Capture location + Weight | 579.42 | 4.70 | 1.00 |
| | Capture location + Sex + Weight | 605.76 | 31.04 | 1.00 |
| Average Number of Stops | Capture location | 48.32 | 0.00 | 0.60 |
| | Capture location + Sex | 50.37 | 2.06 | 0.22 |
| | Capture location + Weight | 51.41 | 3.09 | 0.12 |
| | Capture location + Sex + Weight | 53.16 | 4.84 | 1.00 |

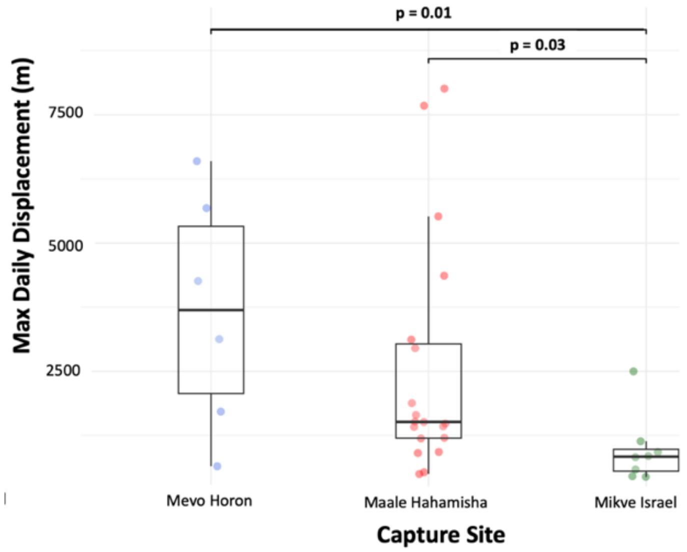


***Figure S4****.* *Max Daily Displacement* among sites ordered by urbanization level: Mevo Horon as our most rural site, Maale Hahamisha semi-rural, Mikve Israel highly urban. Bars indicate significant differences between specific sites.

**Table S3. Effects of site, movement, weight, and sex on alpha diversity and species richness indices**

| Diversity index | Factor | Estimate | SE | p. value |
|---|---|---|---|---|
| Chao1 | Number of Stops | 28.66 | 14.23 | **0.042** |
| | Capture Site - Mikve Israel | 89.65 | 24.89 | **0.0023** |
| | Capture Site - Mevo Horon | 13.46 | 24.94 | 0.596 |
| | Reference (Maale Hahamisha) | 38.55 | 68.93 | 0.583 |
| | Weight | 0.127 | 0.408 | 0.759 |
| | Sex Female | 53.07 | 55.78 | 0.335 |
| | Sex Male | 50.00 | 55.23 | 0.378 |
| Fisher's alpha | Number of Stops | 4.68 | 2.02 | **0.034** |
| | Capture Site – Mikve Israel | 9.98 | 3.52 | **0.007** |
| | Capture Site-Mevo Horon | 0.62 | 4.35 | 0.886 |
| | Reference (Maale Hahamisha) | 0.78 | 9.79 | 0.937 |
| | Weight | 0.01 | 0.05 | .779 |
| | Sex Female | 6.89 | 7.92 | 0.397 |
| | Sex Male | 5.10 | 7.84 | 0.524 |

**Table S4.** Effects of site and movement on alpha diversity and species richness indices after removal of an outlier. note that the main effect of number of stops on the Chao1 index becomes marginally significant, while the Fisher's alpha index effect remains significant.

| **Diversity index** | **Factor** | **Estimate** | **SE** | **p. value** |
|---|---|---|---|---|
| Chao1 | Number of Stops | 31.82 | 18.04 | 0.09 |
| | Capture Site - Mikve Israel | 90.07 | 24.64 | **0.001** |
| | Capture Site-Mevo Horon | 1.72 | 26.61 | 0.94 |
| | Reference (Maale Hahamisha) | 79.68 | 62.91 | **0.22** |
| Fisher's alpha | Number of Stops | 5.56 | 2.54 | **0.04** |
| | Capture Site – Mikve Israel | 8.18 | 3.47 | **0.03** |
| | Capture Site-Mevo Horon | 1.15 | 3.75 | 0.69 |
| | Reference (Maale Hahamisha) | 4.31 | 8.88 | 0.63 |

**Table S5**. Full Pathogen List (see file:"fullpathogenlist")

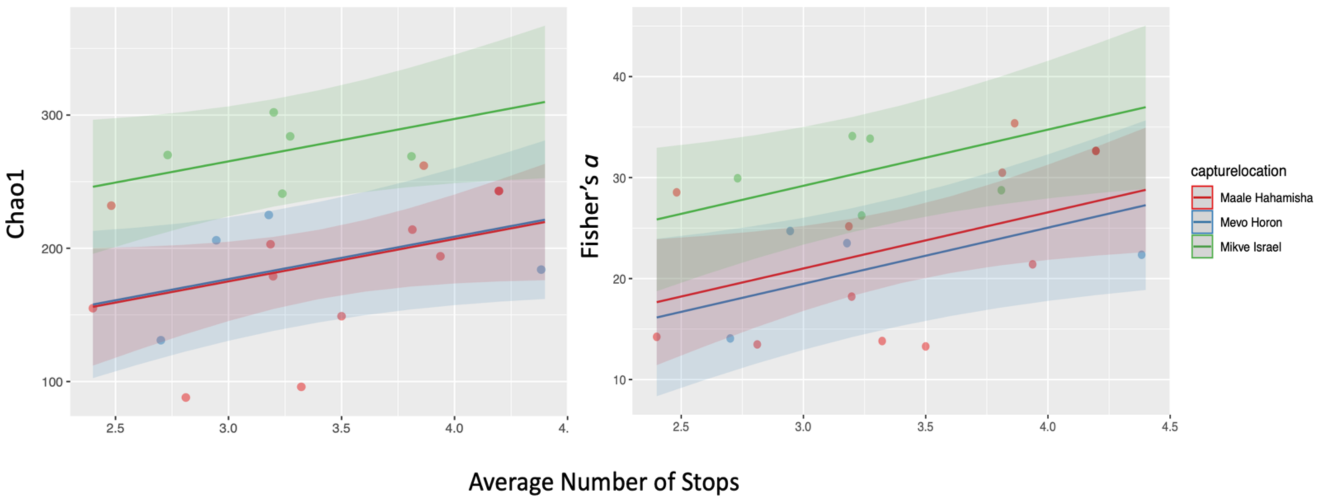


**Figure S4. The effect of pigeon movement on microbiome diversity.** *Results of general linear modeling (GLMM) after the removal of an outlier with very high number of daily stops. Response variables of Chao1 for diversity and Fisher's alpha for evenness, show higher bacteria evenness associated with more daily stops. Shaded areas represent the smoothed 95% confidence interval.*

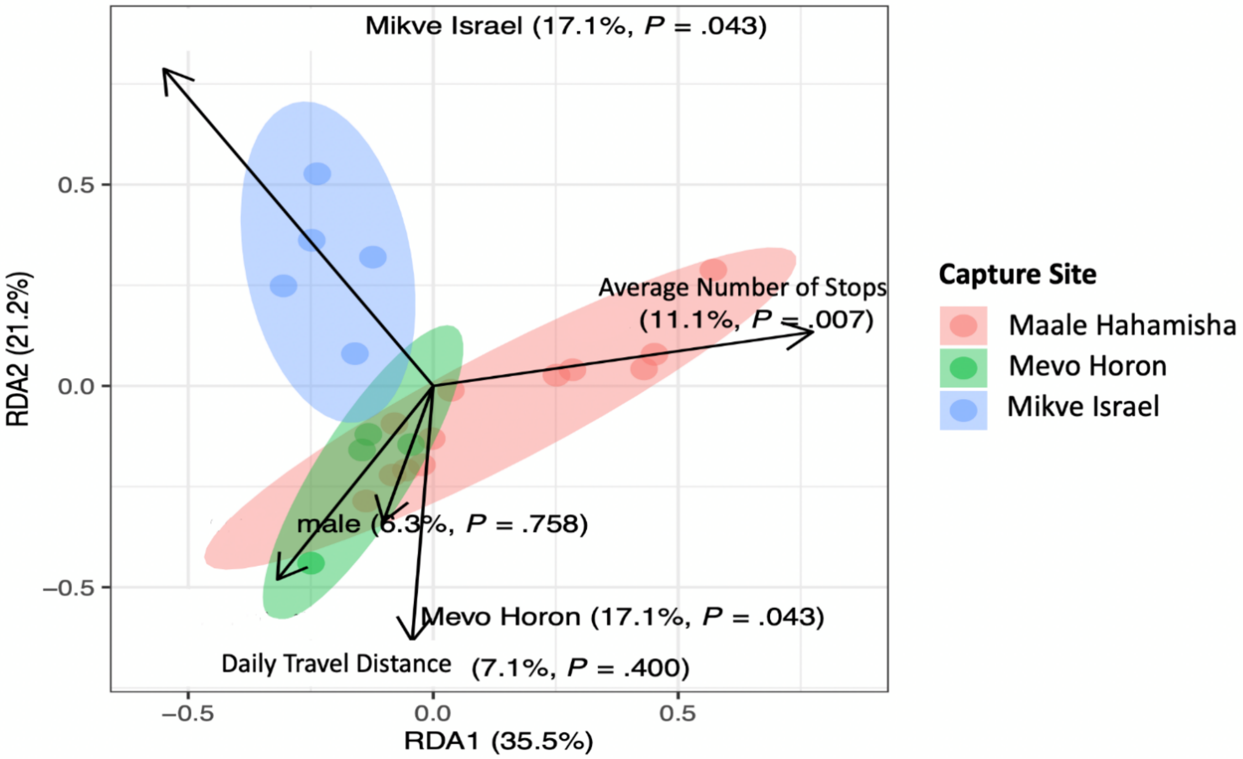


**Figure S6.** *Redundancy Analysis (RDA) of 20 pigeon samples at the genus level. RDA plot shows the result of pigeons grouped capture location. The first and second ordination axes are plotted, explaining 21.2% and 35.5% of the variance. Separation between capture site is significant (P = 0.04, permutation test), in addition to Average Number of Stops (P=0.007).*
